# Bio-based fertilizers shape soil microbiome, resistome and mobilome through metabolism of antibiotic-producing *Streptomyces*

**DOI:** 10.64898/2026.06.29.735163

**Authors:** Taru-Marja Mäkinen, Melina Markkanen, Pietari Lahti-Nuuttila, Kirill Bogdanov, Marko Virta, Jenni Hultman, Johanna Muurinen

## Abstract

*Streptomyces* are abundant soil inhabitants with extensive secondary metabolism and antibiotic resistance traits. Yet, their ecological role in shaping soil antibiotic resistome dynamics remains understudied. Here, we investigated how two different bio-based fertilizers harbouring *Streptomyces* shaped soil resistome and mobilome by combining genome analysis of eight *Streptomyces* isolates to metagenomic profiling of soils before fertilization, within 48 hours after fertilizer application, and six weeks after.

*Streptomyces* genomes showed linkages among antibiotic resistance genes, carbohydrate-active enzymes, and antibiotic-production-associated biosynthetic gene clusters, connecting resistance and biosynthesis to broader metabolic strategies. Relationships between carbon degradation and biosynthesis associated with specific enzyme families, indicating that carbon availability shapes secondary metabolism. We confirmed experimentally that antibacterial potential varied with carbon source, suggesting that microbial activity during manufacturing of the bio-based fertilizers may create localized selection pressures before fertilizers enter the soil.

Fertilization with the studied materials induced modest but consistent shifts in resistome and mobilome without major changes in dominant taxa or overall bacterial abundances, indicating functional reorganization within soil communities. Diversity of antibiotic resistance genes and mobile genetic elements increased, whereas abundance changes were small. Mobile genetic element composition showed stronger responses that were associated with fertilizer inputs, *Streptomyces* abundance, and taxa linked to faecal and resistance sources.

Together, our results show that bio-based fertilizers shape soil resistome primarily through ecological restructuring of resident soil communities, while carbon-dependent microbial activity within fertilizers may enrich resistance. These factors should be considered in manufacturing of bio-based fertilizer as well as in designing agricultural practices.

## Introduction

Antibiotic resistance is an ancient and widespread microbial trait that has existed long before humans discovered antibiotics [1]. Environmental bacteria, particularly antibiotic producers, carry intrinsic antibiotic resistance genes (ARGs) that likely served as ancestral sources of resistance later observed in clinical pathogens [2–4]. These resistance determinants can transfer across distantly related taxa via mobile genetic elements (MGEs) and other horizontal gene transfer mechanisms [5]. Yet, successful transfer depends on ecological context, including taxon co-occurrence, microbial density, and environmental conditions that enable DNA exchange [6,7]. Bio-based fertilizers (BBFs), manufactured from nutrient-rich side-streams such as manure and various compost-based materials, represent environments where these conditions are often met.

Field studies demonstrate that both manure-derived and plant-based amendments can disseminate ARGs and MGEs into soils, altering both their abundance and composition [8,9]. Differences in nutrient regimes, organic matter quality, source streams, and manufacturing practices associated with BBFs can further shape the structure of ARG and MGE pools introduced into or selected within soils [10–13]. Consequently, BBFs may function not only as reservoirs of resistance determinants but also as ecological contexts that influence resistome composition and the structure of associated mobilome pools following fertilization. Here, the resistome refers to the collective repertoire of ARGs within a microbial community, and the mobilome comprises the MGEs that can mediate or reflect the movement and rearrangement of genetic material.

During BBF manufacture, raw materials from nutrient-rich side-streams are gathered and stored in piles and often undergo composting or anaerobic digestion, generating dense, metabolically active microbial communities [14]. Within these communities, members of the genus *Streptomyces* are of particular interest [15]. *Streptomyces* are prolific producers of antibiotics and other bioactive secondary metabolites. This exceptional biosynthetic capacity is supported by large, dynamic genomes enriched in biosynthetic gene clusters (BGCs), and diverse metabolic pathways that enable adaptation to fluctuating environmental conditions [16]. Many BGCs encode intrinsic resistance determinants that prevent self-toxicity, making *Streptomyces* an important environmental reservoir of ARGs [17–19].

Carbon availability is a key regulator of secondary metabolism in *Streptomyces*, with carbon catabolite repression (CCR) influencing both carbon utilization pathways and the expression of BGCs. Different carbon sources have been shown to exert distinct effects on secondary metabolite production [20,21]. Moreover, carbohydrate-active enzymes (CAZymes) are essential to *Streptomyces* ecology, as they enable the degradation of diverse plant- and soil-derived polysaccharides and support the use of a wide range of carbon substrates. Consequently, variation in CAZyme profiles may affect metabolic versatility, influence secondary metabolite production, and shape the intrinsic resistance profiles associated with these metabolic pathways.

In the resource-rich, decomposing and microbial rich environments characteristic of BBFs, sustained antimicrobial production by *Streptomyces* may impose continuous, yet indirect, selective pressures on co-occurring microbial populations. Such pressure could potentially contribute to elevated resistance in the surrounding microbiome and shape the broader resistome and mobilome. Although the roles of secondary metabolism, intrinsic resistance, and horizontal gene transfer in *Streptomyces* are well established [16,20–24], how these processes are shaped by carbon-utilization strategies remains insufficiently understood. In particular, the role of CAZyme-mediated substrate degradation modulating secondary metabolite production and thereby influencing resistance selection dynamics in surrounding microbial communities, remains largely unresolved.

To address these knowledge gaps, we integrated metagenomics of two BBFs and agricultural soils fertilized with them with long-read sequencing of eight *Streptomyces* isolates and formulated three interconnected hypotheses that explore potential links between *Streptomyces* metabolic traits and community-level gene patterns. First, we hypothesize that *Streptomyces* associated with BBFs harbour functionally diverse repertoires of ARGs, BGCs, CAZymes, and MGEs, and their metabolic potential, particularly carbon degradation capacity, is linked to traits involved in antibiotic production and resistance. Second, we hypothesize that a subset of ARGs in *Streptomyces* are associated with MGEs, indicating traces of previous transfer events as well as potential for mobility, and that these ARGs and MGEs are present in the BBFs as well as in fertilized soils. Finally, we hypothesize that BBFs act not only as disseminators of microbial taxa and genes, but also as ecological drivers that influence the structure of soil microbiomes, resistomes, and mobilomes after fertilization. Together, this framework provides a basis for understanding how carbon-mediated metabolism in *Streptomyces* and the use of BBFs may influence microbial community dynamics and resistance gene patterns on short timescales.

## Materials and Methods

### Bio-based fertilizers and sampling

Two bio-based fertilizers (BBFs) were included in this study. The first fertilizer (DSP) was manufactured from pig manure through anaerobic digestion (fermentation) and subsequent separation. The second fertilizer (PLP) was a commercially available product manufactured by Envor Group Oy (Finland), consisting of composted biowaste, food industry by-products, peat, and wood chips. The BBFs were obtained through Natural Resources Institute Finland, where they were sent by the manufacturers.

Soil samples were collected from a field experiment in Jokioinen, Finland (60°48′15.6"N 23°27′06.5"E) with no organic fertilization in the previous five years. The field trial used a randomized complete block design with four replicate plots (≥120 m²) per BBF treatment (8 plots total). BBFs were surface-applied and incorporated into the top 10 cm within 24 hours. Topsoil samples, from 0 to 20 cm depth, were collected at three timepoints: one day before fertilization (before), within 48 hours after fertilization (=after), and six weeks after the fertilization (=six weeks after). Sampling was performed by collecting soil from six spots per plot into a cleaned bucket where the soil was mixed and transferred into a sterile plastic bag. Samples were transported in coolers with cooling elements to the laboratory on the same day and stored at 8°C until processing (within two days). All samples were sieved through a 5 mm screen, mixed, and preserved at −20°C for DNA extraction. The abbreviations, treatments, and treatment times for fertilizers and soil samples are listed in Supplementary Table S1, sheet 1.

### Bacterial isolation, strains, and growth conditions

Bacteria were isolated from DSP and PLP by suspending 0.5 g of each in 50 ml of 1X PBS, followed by serial dilutions (10⁻² to 10⁻⁴) using 1% saline. Aliquots (50 μL) were spread onto Gause’s No. 1 and International *Streptomyces* project Medium No. 7 (ISP7; without glycerol) (HiMedia Laboratories, Mumbai, India) [25] supplemented with Nystatin (10,000 U/ml, Gibco, West Sussex, UK) and Nalidixic acid (30 μg/ml, Sigma-Aldrich, St. Louis, MO, USA) in a 1:1000 ratio. Gause’s No. 1 medium composition was (g/L): soluble starch, 20; NaCl, 0.5; FeSO_4_·7H_2_O, 0.01; K2HPO_4_·3H_2_O, 0.64; KNO_3_, 1; MgSO_4_·7H_2_O, 0.5; agar, 14–15. Plates were incubated at 28°C for four days. Pure cultures were obtained by restreaking single colonies and incubating as above. Eight isolates displaying *Streptomyces*-like morphology (Supplementary Fig. S1) were stored at 4°C and at -70°C as glycerol stocks. For DNA extraction, isolates were cultured on agar plates at 28°C for 3-4 days until sufficient biomass was obtained.

### Antimicrobial activity experiment

Antimicrobial activity of selected isolates (BBF42_03, BBF42_NS, BBF45_04, BBF45_11) was assessed using a modified cross-streak assay [26]. *Kitasatospora aureofaciens* (prev. *Streptomyces*) strain H-51 was included as a positive control, and *Escherichia coli* DH5α served as the indicator strain. Isolates were grown on ISP media (ISP4, ISP5) at 28°C for one week prior to perpendicular streaking of the indicator strain. Plates were incubated at 37°C, and antimicrobial activity was evaluated based on inhibition of *E. coli* growth. Detailed assay conditions and information on the control strain are provided in Appendix S1 and results Supplementary Fig. S3.

### DNA extraction and sequencing

DNA from fertilizers, soils, and bacterial isolates was extracted using the DNeasy PowerLyzer PowerSoil Kit (Qiagen, Hilden, Germany) with minor modifications. Weighed samples and harvested biomass from bacterial isolates were transferred to PowerBead tubes containing 600 μl of PowerBead Solution and stored overnight at 4°C. Cell lysis was performed using a horizontal vortex adapter for 15 minutes. An additional 15 min incubation at 4°C was included during inhibitor removal steps. DNA was eluted in 50 μl of elution buffer, incubated for two minutes at room temperature, and collected by centrifugation. The concentration and quality of the extracted DNA were determined with the NanoDrop One spectrophotometer (Thermo Fisher Scientific, Waltham, MA, USA). The whole genome sequencing was performed using the PacBio Sequel II system (Pacific Biosciences, Menlo Park, CA, USA) with SMRTbell library at the Institute of Biotechnology, University of Helsinki. Shotgun metagenomic sequencing was performed using Illumina NovaSeq6000 platform (Illumina, San Diego, CA, USA) with 2 x 150 bp paired-end reads and Nextera XT library preparation (Novogene, Cambridge, UK).

### Bioinformatics for metagenomic data

Raw paired-end reads were filtered and trimmed using Cutadapt v3.5 [27] (adapters, q < 20; length < 50 bp). Read quality was assessed with FastQC v.0.11.9 [28] and MultiQC v.1.19 [29]. Microbial community composition was characterized using phyloFlash v3.4.2 [30], against SILVA 138.1 SSU rRNA reference database [31]. Taxonomic names were updated to match Genome Taxonomy Database (GTDB) version R226 [32] nomenclature and current classification standards for the figures. Antibiotic resistance genes (ARGs) were identified by mapping quality-filtered reads to ResFinder database v2.6.0 [33] using Bowtie2 v2.5.3 [34] with parameters: -D 20 -R 3 -N 1 -L 20 -i S,1,0.50. ResFinder database was dereplicated at 99% sequence identity using CD-HIT v4.8.1 [35] prior to mapping. Mobile genetic elements (MGEs) were detected using the same mapping approach against the MobileGeneticElementDatabase v1.1 [36]. Mapped reads were processed with SAMtools v1.21 [37], with paired and singleton mappings counted as single hits. The abundance of *Streptomyces* isolates in the metagenomes was quantified by mapping the quality-filtered reads to isolate assemblies using Bowtie2 v2.5.3 with identical parameters.

### Bioinformatics for genomic data

PacBio HiFi reads were adapter-trimmed with HifiAdapterFilt v2.0.0 [38], and quality-checked with FastQC v.0.11.9 and MultiQC v.1.19. De novo assemblies were generated using Hifiasm v0.19.5 [39,40], selecting the primary contig graph (‘prefix’.bp.p_ctg.gfa) for downstream analysis. Assembly quality was evaluated using Quast v5.3.0 [41]; completeness and contamination with CheckM2 v1.1.0 [42]. Reads were mapped back to assemblies with minimap2 v2.24 [43], and contigs with <10x were removed, reducing contamination while maintaining ≥97% completeness.

Taxonomic assignment was performed using GTDB-Tk v2.5.2 [44] against GTDB R226. Closest reference genomes were retrieved from NCBI RefSeq/Genbank for comparative analyses (Supplementary Table S5, sheet 1). *Streptomyces coelicolor* A3(2) (GCF_000203835.1) was included as model reference in comparative analyses. Pangenome analysis with Roary v3.13.0 (90% BLASTP identity, 99% clustering threshold) [45], and maximum likelihood phylogeny with RAxML v8.2.12 (GTR+GAMMA, 200 bootstrap) [46] were used to assess genome similarity.

Protein-coding sequences were predicted and annotated using Bakta v1.11.0 [47] with the full database v6.0 [48]. Carbohydrate-active enzymes (CAZymes) were identified with dbCAN3 v5.1.2 [49], against CAZy database v5.2 (Carbohydrate-Active Enzymes database, http://www.cazy.org/) [50], retaining only high-confidence annotations (“Recommended Results”). ARGs and MGEs were detected using BLASTN v2.16.0 [51] against the dereplicated ResFinder [33] and MobileGeneticElementDatabase [36] with e-value ≤ 0.01. Secondary metabolite biosynthetic gene clusters (BGCs) were predicted using antiSMASH v8.0.4 [52], and cluster annotations were parsed using custom Python script.

### Statistical analyses of the metagenomes

Analyses were conducted in R Statistical Software v4.5.0 [53] within RStudio v2025.5.0.496 [54], using tidyverse 2.0.0 [55] and vegan v2.7-2 [56]. Normality was assessed via Shapiro-Wilk tests (fertilizer group excluded, n = 2); abundance data were analyzed non-parametrically regardless of test outcome given small group sizes (n = 4) and the inherently non-negative, right-skewed nature of metagenomic count data. Differences between treatments and the soil before were assessed using unpaired Welch’s t-tests (Shannon diversity) and unpaired Wilcoxon rank-sum tests (abundances). Community composition was analyzed using Bray-Curtis dissimilarities, PERMANOVA (adonis2, 999 permutations), and tests for homogeneity of dispersions (betadisper, permutest). Benjamini–Hochberg (BH) correction was applied across to all univariate tests and pairwise PERMANOVA.

Bacterial 16S rRNA gene profiles were processed with phyloseq v1.52.0 [57], excluding mitochondrial and chloroplast reads. Alpha diversity was estimated by Shannon index (raw bacterial counts), and beta diversity using Bray-Curtis dissimilarities of genus-level total-sum-scaled (TSS) abundances. Ordination was performed using Principal Coordinates Analysis (PCoA) and visualized both including and excluding fertilizer samples, with PERMANOVA applied globally and pairwise against soil before. Taxonomic composition was visualized at order level using GTDB R226 nomenclature.

ARG and MGE read counts were normalized based on gene length and the sample-specific bacterial 16S rRNA gene read counts, and by multiplying them by the length of reference 16S rRNA gene (1,541 bp), providing a proxy for gene abundance relative to microbial load. Diversity was assessed using Shannon index, and community composition by Nonmetric Multidimensional Scaling (NMDS) on Bray-Curtis dissimilarities of Hellinger-transformed data to reduce the influence of highly abundant genes. Significant gene contributors were identified with envfit (999 permutations, BH-corrected p-values < 0.05, top 10 vectors by r² displayed). MGEs significantly associated with ordination space were annotated using BLASTN v2.16.0 against NCBI nt database (January 2020; e-value ≤ 1×10⁻¹⁰; up to 5 non-redundant hits per MGE retained by bit score and percent identity, Supplementary Table S1, sheet 5).

Relative abundances of *Streptomyces* and selected bacterial genera associated with faecal inputs and antimicrobial resistance (*Enterococcus, Staphylococcus, Klebsiella, Acinetobacter, Enterobacter, Escherichia–Shigella, Campylobacter, Arcobacter, Moraxella, Serratia, Aeromonas, Providencia, Salmonella, Bacteroides, Vibrio, and Streptococcus*) [58,59]; *Pseudomonas* excluded due to broad environmental distribution) were overlaid onto ARG and MGE ordinations as nonlinear surfaces using ordisurf (GAM, k = 5, shrinkage penalty; n = 22). MGE abundance was additionally overlaid onto the ARG ordination and vice versa. ARG, MGE, and total bacterial Shannon diversity were fitted as linear envfit vectors (999 permutations; p < 0.05 shown). A combined *Streptomyces* metric summed 16S rRNA relative abundance and length- and 16S-normalized isolate contig abundance. Treatment effects on focal taxa were tested with Wilcoxon rank-sum tests.

### Functional gene analysis of the isolates

ARG and MGE hits were filtered by bitscore and query coverage thresholds (ARGs: bit-score ≥100, query coverage ≥50%; MGEs: bit-score ≥50, query coverage ≥30%) and overlapping hits on the same contig were resolved by retaining the highest-scoring alignment. CAZyme annotations were curated to the family level, with multi-domain proteins split and counted per family. BGC predictions were linked to MIBiG database v4.0, retaining matches with ≥15% cluster similarity; BGCs with predicted antibiotic bioactivity, as defined by MIBiG bioactivity terms, were used in downstream analyses except where otherwise indicated. Filtered profiles were compiled into presence/absence matrices (ARGs, antibiotic BGCs) and count matrices (CAZyme families, MGE types) and visualized alongside a maximum-likelihood phylogeny as layered heatmaps (Supplementary Table S5, sheets 2–9). ARGs and MGEs identified in the isolates were screened against the metagenomic datasets to assess their occurrence across treatment groups.

Pairwise compositional dissimilarity was assessed using Mantel tests (Jaccard distances, 999 permutations). Cross-category associations were assessed by Spearman rank correlations between pairwise Jaccard similarities and between CAZyme class counts and BGC product type counts (all predicted BGCs), with BH-corrected p-values. Genomic co-localization of ARG-MGE and ARG-antibiotic-production-associated BGC pairs was assessed by pairwise distances on the same contig using BLASTN-derived coordinates (ARGs, MGEs) and antiSMASH-defined BGC boundaries, with co-localization defined as ≤ 10 kb separation.

## Results

### Fertilization reshapes bacterial community structure with minimal effects on dominant phyla and abundance

Across the 22 metagenomes, sequencing generated 32.2 ± 3.4 million paired-end reads (mean ± SD), of which 0.052 ± 0.016% were identified as 16S rRNA reads and 95.6 ± 2.6% were classified as bacterial (Supplementary Table S1, sheet 1). Fertilizers (DSP, PLP) and soils (before, after, and six weeks after) exhibited distinct bacterial community profiles. Fertilizers were enriched in *Bacteroidota* and *Pseudomonadota*, whereas soils were characterized by *Pseudomonadota*, *Actinomycetota*, and *Verrucomicrobiota* (Fig. 1A, Supplementary Table S1, sheet 2).

**Figure 1.**
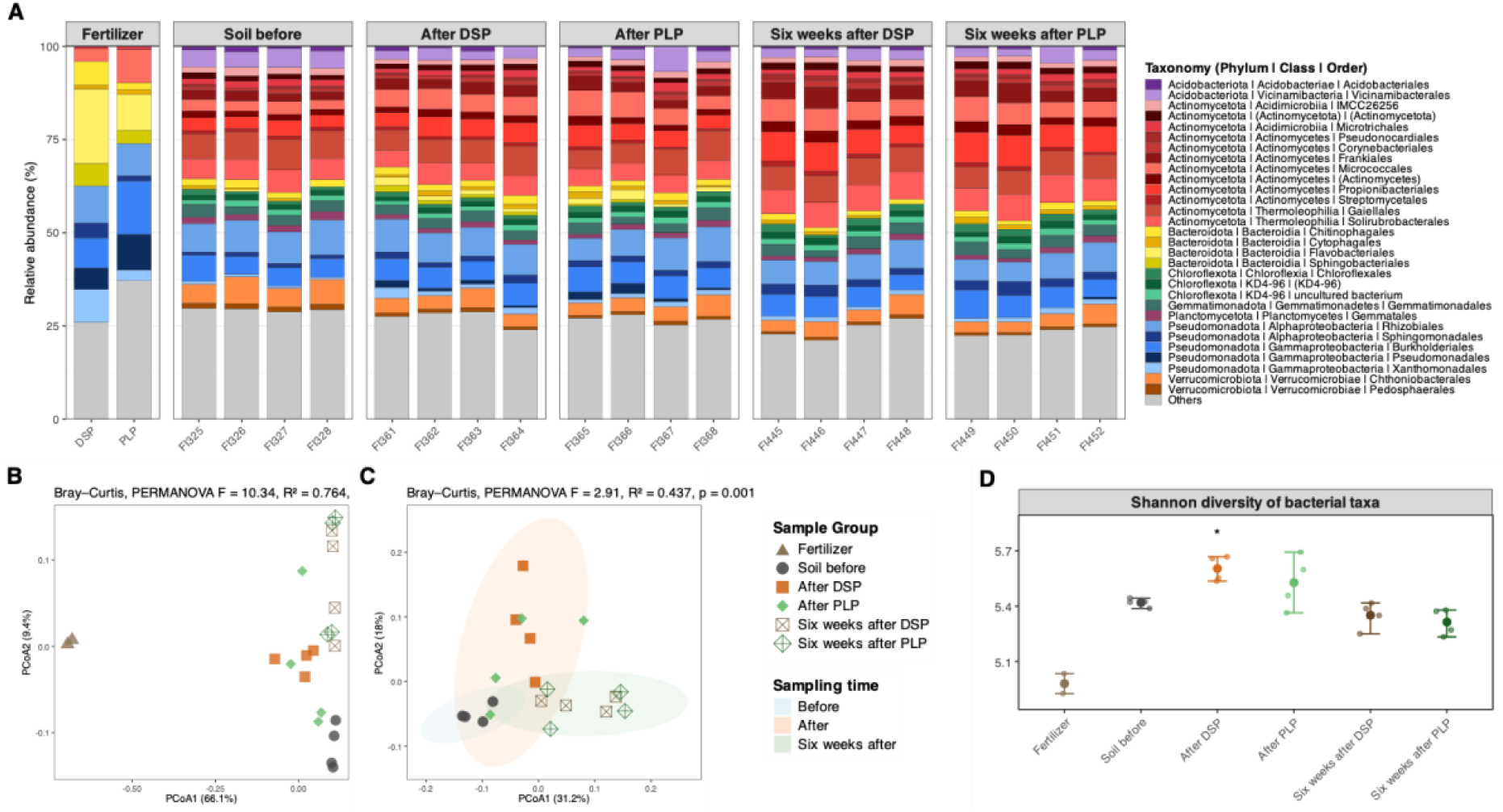
Taxonomic composition, community structure, and diversity of bacterial communities across sample groups. Relative abundance of the 30 most abundant bacterial orders (≥ 0.5% relative abundance shown) following total-sum-scaling (TSS) within bacterial reads (A). All remaining orders are grouped as “Others”. Principal coordinate analysis (PCoA) of genus-level bacterial community composition based on Bray–Curtis dissimilarities, shown for all samples including fertilizers (B). PcoA of soil samples only; shaded ellipses represent 95% confidence regions (t-distribution) for three sampling time points: before, after, and six weeks after (C). Each point represents an individual sample, with colours and shapes indicating sample group; percentages on axes indicate the proportion of variation explained by each axis. PERMANOVA statistics are displayed within the corresponding panels. Shannon diversity of bacterial communities across sample groups (D); individual points represent samples; horizontal points group means, and error bars represent the observed minimum-maximum range. Asterisks indicate significant difference relative to the soil before group based on unpaired Welch’s t-tests with Benjamini–Hochberg correction (*p(BH) < 0.05; Supplementary Table S3). Sample sizes were n = 2 for fertilizer and n = 4 for all other groups.

Genus-level PCoA showed a clear separation between the fertilizers and soil sample groups (PERMANOVA, R^2^ = 0.76, F = 10.34, p = 0.001; Fig. 1B), while soil samples alone displayed subtler but significant clustering by sampling time (PERMANOVA, R^2^ = 0.44, F = 2.91, p = 0.001; Fig. 1C). Pairwise PERMANOVA confirmed that all fertilized soils differed from the soil before (p(BH) = 0.022 for all comparisons; Supplementary Table S2). Group dispersion differed when fertilizers were included (PERMDISP, F = 13.57, p = 0.001), indicating that differences in both centroid location and dispersion contributed to the observed separation. In contrast, dispersion did not differ significantly among soil samples (PERMDISP, F = 2.50, p = 0.11), suggesting that soil-only clustering was primarily driven by differences in community composition. Despite these compositional shifts, dominant phyla remained stable and overall bacterial abundance did not change (Wilcoxon tests, all p(BH) ≥ 0.15; Fig. 1A; Supplementary Table S3). Focal taxa abundances (including *Streptomyces* and selected genera associated with faecal inputs and ARGs) likewise did not differ among treatments (all p(BH) > 0.12; Supplementary Table S3). Shannon diversity increased immediately after DSP (Welch’s t-test, p(BH) = 0.036), while other comparisons were non-significant (p(BH) ≥ 0.115; Fig. 1D, Supplementary Table S3).

### Fertilization increases resistome and mobilome diversities but only slightly increases abundances of ARGs and MGEs

Across metagenomes, we detected a broad collection of ARG and MGE variants (Supplementary Table S1, sheets 3,4). Fertilizers exhibited the highest ARG and MGE abundance and diversity (Fig. 2A–D). In soils, fertilization primarily increased diversity, with modest effects on abundance (Supplementary Table S3). ARG diversity increased after both fertilizers and remained elevated six weeks after DSP but not PLP (Welch’s t-test, p(BH) = 0.008; Fig. 2B). Similarly, MGE diversity increased after both fertilizers and remained above the soil before levels at six weeks (p(BH) ≤ 0.012; Fig. 2D). ARG abundance showed small but significant increases after fertilization, except six weeks after PLP (Wilcoxon tests, p(BH) = 0.041; Fig. 2A), whereas MGE abundance did not significantly differ among soil groups (p(BH) ≥ 0.062; Fig. 2C).

**Figure 2.**
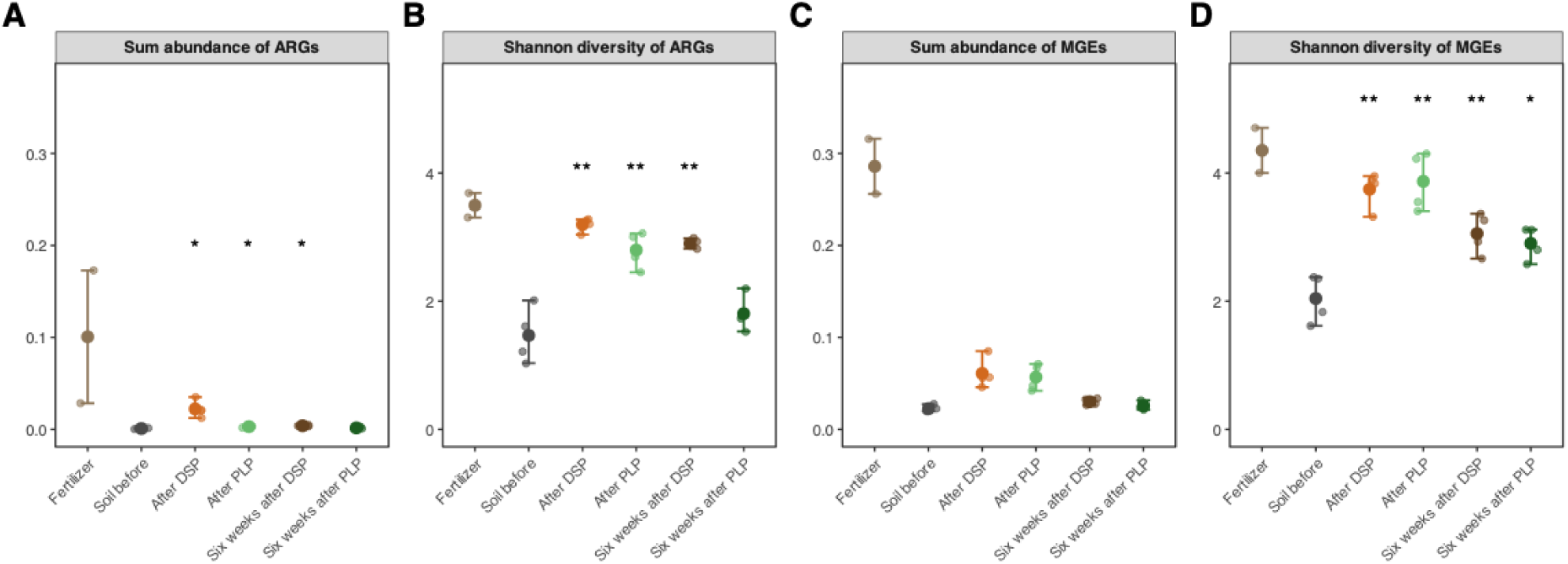
Abundance and diversity of antibiotic resistance genes (ARGs) and mobile genetic elements (MGEs) across sample groups. Total normalized abundance (A, C) and Shannon diversity (B, D) of ARGs (A, B) and MGEs (C, D) are shown for each sample group. Abundances were normalized to gene length and bacterial 16S rRNA gene counts (see Methods); individual points represent samples; horizontal points group means, and error bars represent the observed minimum-maximum range. Asterisks indicate significant differences relative to the soil before group based on unpaired Wilcoxon rank-sum tests for abundance and unpaired Welch’s t-tests for diversity with Benjamini–Hochberg correction (*p(BH) < 0.05, **p(BH) < 0.01; Supplementary Table S3). Sample sizes were n = 2 for fertilizer and n = 4 for all other groups.

### Fertilization restructures resistome and mobilome composition

Non-metric multidimensional scaling (NMDS) ordinations revealed significant shifts in ARG and MGE compositions along a gradient from the soil before to fertilizers (PERMANOVA, ARG: R^2^ = 0.48, F = 2.98; MGE: R^2^ = 0.57, F = 4.17; p = 0.001; Fig. 3A, E). Dispersion patterns differed between datasets: MGE profiles showed homogeneous dispersion (PERMDISP, F = 0.77, p = 0.581), indicating compositional shifts, whereas ARG profiles exhibited increased dispersion (PERMDISP, F = 6.90, p = 0.001), suggesting both compositional change and higher variability. MGEs showed clearer separation than ARGs, forming distinct clusters between fertilizers and the soil before, whereas ARGs displayed a more continuous distribution across treatments. Over time, resistomes and mobilomes in fertilized samples shifted partially toward the soil before composition, with more constituently for MGEs than ARGs (Fig. 2; Fig. 3A, E).

**Figure 3.**
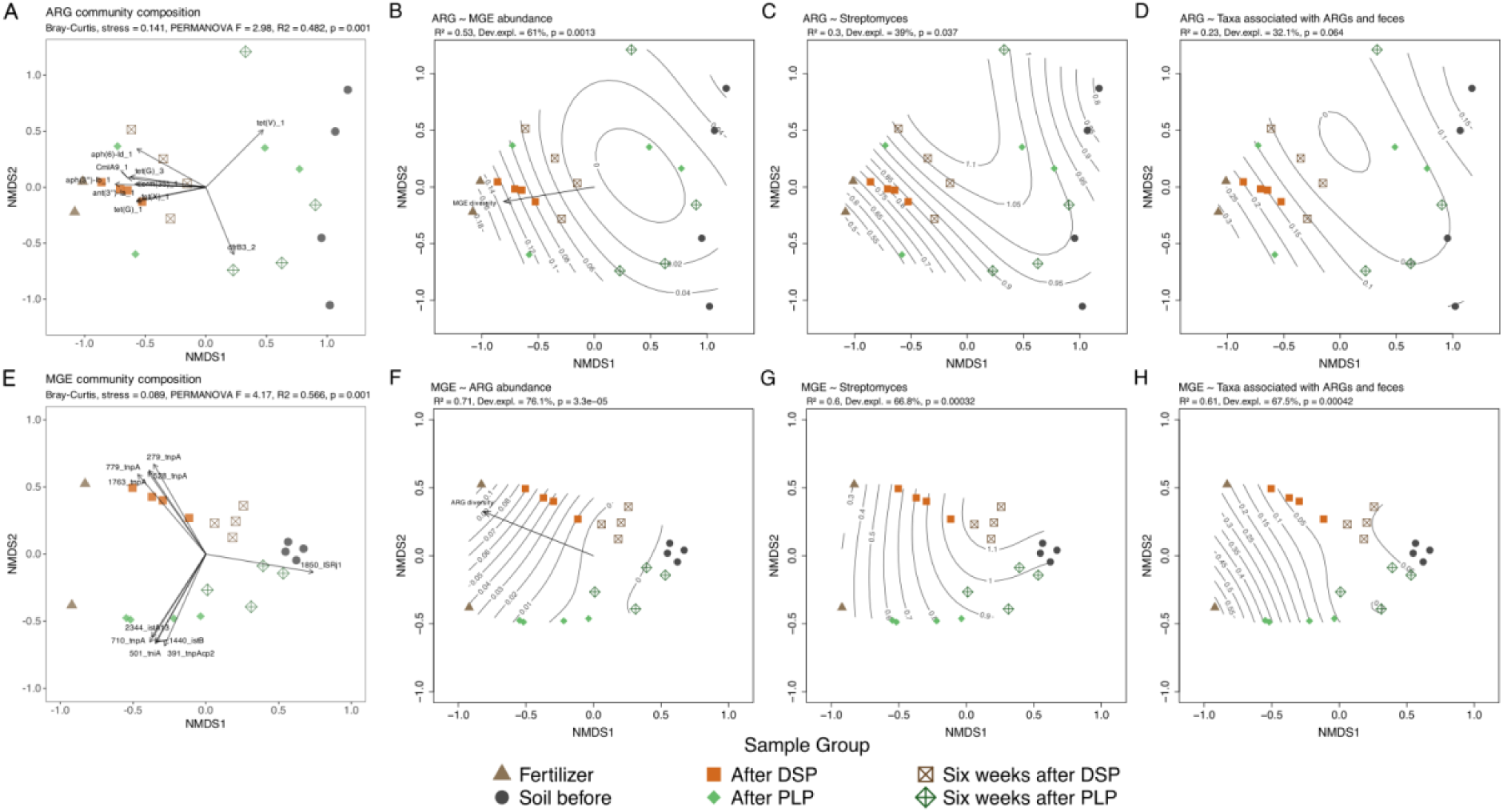
Community structure of antibiotic resistance genes (ARGs) and mobile genetic elements (MGEs). Non-metric multidimensional scaling (NMDS) ordinations of ARG (A) and MGE (E) community composition based on Bray–Curtis dissimilarities of Hellinger-transformed abundances. Points represent individual samples coloured and shaped by treatment group. Arrows indicate the 10 gene features most strongly associated with ordination space (envfit, 999 permutations; Supplementary Table S4), with arrow length proportional to association strength (R²); stress values and PERMANOVA statistics are shown within panels. Generalized additive model surfaces (ordisurf, GAM k = 5) illustrate the association of ARG community composition with MGE abundance (B), *Streptomyces* abundance (C), and abundance of faecal- and ARG-associated taxa (D), and of MGE community composition with ARG abundance (F), *Streptomyces* abundance (G), and faecal- and ARG-associated taxa (H). Where significant (p < 0.05, 999 permutations), Shannon diversity of MGEs (B) and ARGs (F) are shown as linear vectors. Contours represent fitted values with model fit statistics (R², deviance explained, p shown in each panel).

For ARGs, samples formed a continuum from the soil before to fertilizers. DSP-fertilized soils clustered near fertilizers and remained relatively stable, whereas PLP-fertilized soils were more dispersed and partially overlapped with the soil before samples (Fig. 3A). Multiple ARGs strongly aligned with ordination axes (R² = 0.50–0.87; Supplementary Table S4), including aminoglycoside, tetracycline, and MLSB resistance genes, which were linked to fertilized soils. In contrast, one tetracycline resistance gene (*tet(V)_1*) and trimethoprim resistance gene (*dfrB3_2*) were associated with the soil before and a subset of PLP-fertilized samples.

MGE profiles showed a clearer separation, with fertilizers and fertilized soils aligned with multiple transposase genes (R² = 0.81–0.90; Supplementary Table S4), while the soil before samples were strongly associated with the insertion sequence IS*Rj1* (R² = 0.88; Fig. 3E). BLAST analysis indicated that IS*Rj1* was predominantly associated with *Bradyrhizobium* species (Supplementary Table S1, sheet 5). Transposase genes associated with DSP-fertilized soils showed closest matches to plasmids and genomes from clinically relevant or opportunistic genera, including *Enterobacter*, *Escherichia coli*, *Klebsiella*, *Salmonella*, and *Acinetobacter*. PLP-fertilized soils were associated with broader set of MGEs, with the closest matches spanning both environmental taxa (e.g. *Pseudomonas*, *Stutzerimonas*) and clinically relevant and opportunistic genera (*Citrobacter*, *Serratia*, *Stenotrophomonas*).

ARG composition aligned with gradients in MGE abundance (R² = 0.53, dev.expl. = 61%, p = 0.001; Fig. 3B), *Streptomyces* abundance (R² = 0.30, dev.expl. = 39%, p = 0.037; Fig. 3C), and abundance of faecal- and ARG-associated taxa (R² = 0.23, dev.expl. = 32%, p = 0.064; Fig. 3D), although the latter was not significant. MGE composition showed stronger associations, with ARG abundance (R² = 0.71, dev.expl. = 76%, p < 0.001; Fig. 3F), *Streptomyces* abundance (R² = 0.60, dev.expl. = 67%, p < 0.001; Fig. 3G), and abundance of faecal- and ARG-associated taxa (R² = 0.61, dev.expl. = 68%, p < 0.001; Fig. 3H) explaining substantial variation in ordination space. This tight coupling was further supported by a significant linear association between MGE Shannon diversity and ARG ordination space (envfit R² = 0.67, p = 0.001; Fig. 3B), and between ARG Shannon diversity and MGE ordination space (R² = 0.78, p = 0.001; Fig. 3F). Increasing *Streptomyces* abundance corresponded to a shift toward the soil before resistome structure, whereas lower abundance aligned with fertilizer-associated profiles.

### High-quality *Streptomyces* genome assemblies reveal multiple species

Genome sequencing produced high-quality assemblies for all isolates following the coverage-based filtering (Supplementary Table S5, sheet 1). Final assemblies ranged from 7.27 to 8.45 Mb, contained one to six contigs per genome, and had GC contents of 71.9–73.3%. Genome completeness was 98.0–100% with ≤1.5% contamination. Across genomes, 6,111–7,492 coding sequences were predicted with an average coding density of 87.7%. Of these, 5.34–10.0% encoded hypothetical proteins with no assigned functions. Taxonomic assignment placed all isolates within *Streptomyces*, representing five species. Two isolates (BBF42_03, BBF42_BW) were assigned to *S. speibonae*, three isolates (BBF45_07, BBF45_11, BBF45_15) to *S. cellulosae* and BBF42_NS to *S. diastaticus* (Supplementary Table S5, sheet 1). Two isolates (BBF45_04, BBF45_12) were closely related (99.5% average nucleotide identity (ANI)) to *Streptomyces* sp010548465, with *S. viridosporus* (GCF_002078235.1; 88.2% ANI) as the closest named species.

### Functional gene profiles are lineage-specific

Functional gene profiles mirrored phylogeny across ARGs, antibiotic-production-associated BGCs, CAZymes, and MGEs, while lineage-specific differences were observed (Fig. 4, Appendix S2, Supplementary Table S5, sheets 2–9). Resistance genes spanned several antibiotic classes, including aminoglycosides, β-lactams, and macrolides. The BGC and CAZyme repertoires also had distinct, lineage-specific compositions and a small, conserved core (e.g. albaflavenone biosynthesis). Notably, *S. diastaticus* lacked polysaccharide lyases (PL) and showed limited glycoside hydrolase (GH) diversity. Among MGEs, only IS*Cau1* and different variants of *tnpA* were detected. To assess environmental relevance, isolate-associated ARGs and MGEs were searched against metagenomes. Only three genes (*ole*(C), 1308_tnpA, 2212_tnpA) were detected, occurring at low and inconsistent abundance across treatments or time points (Supplementary Table S1, sheet 1).

**Figure 4.**
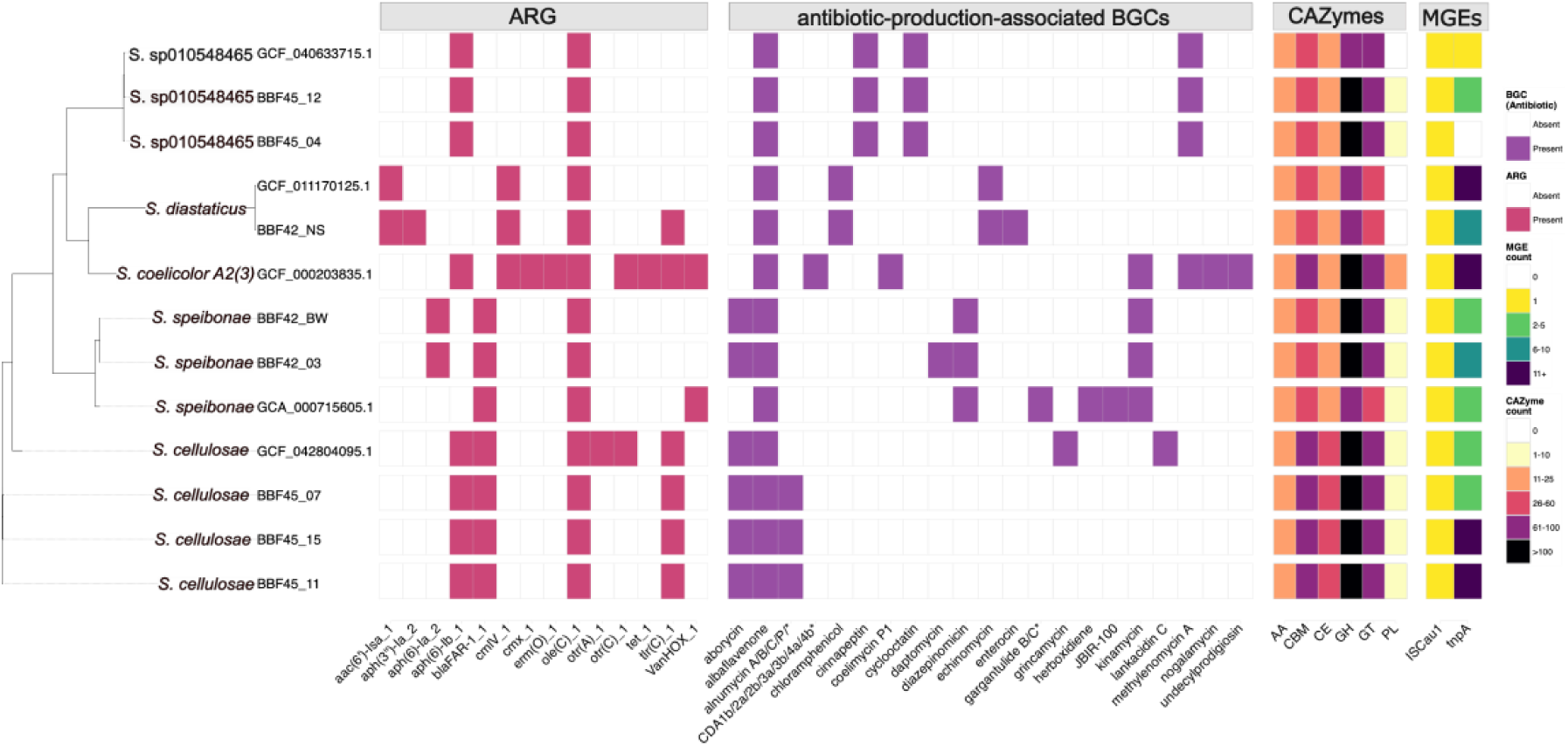
Distribution of functional genes across *Streptomyces* genomes. Functional gene profiles for antibiotic resistance genes (ARGs), antibiotic-production-associated biosynthetic gene clusters (BGCs), carbohydrate-active enzyme (CAZyme) families, and mobile genetic elements (MGEs) are shown across 13 *Streptomyces* genomes. A maximum-likelihood phylogeny is displayed on the left, with gene content visualized as aligned heatmaps. Presence–absence data are shown for ARGs and BGCs, and counts are shown for CAZyme families and MGE classes. CAZyme classes are grouped as AA (auxiliary activities), CBM (carbohydrate-binding modules), CE (carbohydrate esterases), GH (glycoside hydrolases), GT (glycosyltransferases), and PL (polysaccharide lyases). Colour scales indicate presence–absence or abundance, as shown in the legends.

### Functional gene categories co-vary but show limited linkage to mobile genetic elements

Mantel tests showed positive associations among all four functional gene categories (all p(BH) ≤ 0.048), with the strongest associations between ARG and BGC profiles (Mantel r = 0.87), followed by CAZyme–BGC (r = 0.83), and CAZyme–ARG (r = 0.82). In contrast, MGE profiles showed weaker associations with the other categories (r = 0.25–0.40). Consistent patterns were observed in isolate-level Jaccard similarities (Fig. 5), where similarities in ARG, CAZyme, and antibiotic-production-associated BGC profiles co-varied positively, with the strongest correspondence between BGC and CAZyme similarity (Fig. 5C). In line with the weak ARG–MGE associations, MGE similarity showed limited alignment with ARG and BGC patterns (Fig. 5 D, F), and no ARG–MGE co-localization was detected within isolate genomes. Instead, the *ole*(C) resistance gene was found embedded within antibiotic-production-associated BGCs—specifically echinomycin or cinnapeptin clusters—in three isolates (BBF42_NS, BBF45_04, BBF45_12).

**Figure 5.**
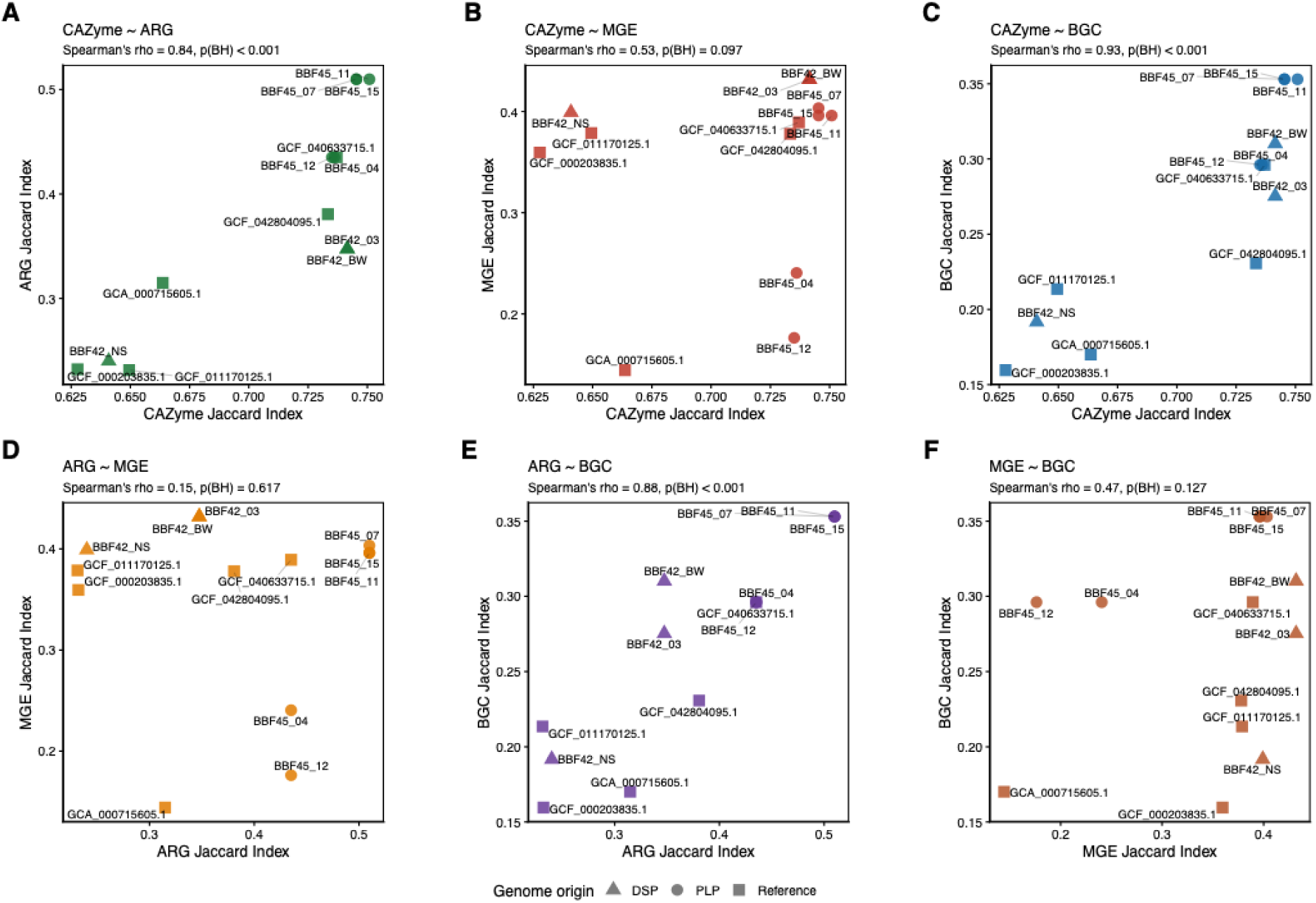
Associations among functional gene repertoires across *Streptomyces* genomes. Spearman rank correlations of pairwise Jaccard similarity indices for antibiotic resistance genes (ARGs), carbohydrate-active enzymes (CAZymes), antibiotic-production-associated biosynthetic gene clusters (BGCs), and mobile genetic elements (MGEs) across 13 *Streptomyces* genomes. Panels show: (A) CAZyme ∼ ARG, (B) CAZyme ∼ MGE, (C) CAZyme ∼ BGC, (D) ARG ∼ MGE, (E) ARG ∼ BGC, and (F) MGE ∼ BGC. Each point represents an individual genome, with symbols indicating genome origin: triangles (DSP-derived isolates, circles (PLP-derived isolates), and squares (reference genomes). Spearman’s rho and associated Benjamini–Hochberg corrected p-values (p(BH)) are shown in each panel subtitle.

To further examine the relationship between carbon degradation capacity and biosynthetic potential, Spearman rank correlations were calculated between CAZyme class abundances and BGC product type counts across all predicted regions (Supplementary Fig. S2). Type II polyketide synthase regions showed strong positive associations with total CAZyme count and carbohydrate-binding module, carbohydrate esterase, and glycoside hydrolase abundances (ρ = 0.82–0.89, p(BH) ≤ 0.05), while non-ribosomal peptide synthetase regions showed consistent inverse relationships across the same families (ρ = −0.81 to −0.90, p(BH) ≤ 0.05).

To assess whether the genomic correspondence between CAZyme and BGC repertoires (Fig. 5C) was reflected in phenotypic antibacterial activity, a qualitative cross-streak inhibition assay against *Escherichia coli* was performed (Appendix S1). All tested isolates exhibited inhibition against *E. coli*, with generally stronger inhibition observed on starch-based ISP4 compared to glycerol-based ISP5 media (Supplementary Fig. S3). The control strain, *Kitasatospora aureofaciens* H-51, showed complete inhibition on ISP4 and partial inhibition on ISP5.

## Discussion

Our findings show that bio-based fertilizers (BBFs) and *Streptomycetes* communities can influence soil resistome dynamics through ecological processes that begin before the fertilizers are applied to soil. During manufacture and storage, bacterial communities in BBFs remain active, allowing them to develop substrate-specific functions, with *Streptomyces* playing a key role in secondary metabolite production and carbon cycling [15]. Consequently, BBFs enter soil with established microbial community structures and functions.

Our eight isolates contained a wide variety of carbohydrate-active enzymes (CAZymes) and biosynthetic gene clusters (BGCs), and the strong similarity in their composition suggests that these functions may be interconnected rather than independent. The relationship between carbon degradation capacity and metabolite potential depended on the specific CAZyme families and BGC types, indicating that carbon availability affects biosynthetic pathways in distinct ways rather than increasing biosynthetic diversity uniformly. Isolates with more diverse CAZyme profiles may therefore maintain the carbon-derived signals that modulate antibiotic production and antagonistic interactions [16,20,21], which aligns with the phenotypically observed carbon-dependent antibacterial activity. Together, the sustained metabolite production during storage may impose localized selection pressure of co-occurring microorganisms, thereby shaping resistance-related traits before fertilization.

Resistance in *Streptomyces* appeared closely tied to broader functional traits. CAZymes, antibiotic-production-associated BGCs, and resistance genes formed phylogenetically structured associations, while connections to mobile genetic elements (MGEs) were weak. In contrast to our hypothesis, no ARG-MGE co-localization was detected. This suggests that resistance in *Streptomyces* is built into the intrinsic metabolic and ecological strategies rather than acquired via horizontal gene transfer, which aligns with how *Streptomyces* genomes evolve primarily through duplication and rearrangement of BGCs [16,17,19,22,23]. In addition, only a subset of the isolate-derived ARGs and MGEs were detected in metagenomes, typically at low abundances. This pattern is consistent with the idea that soil resistomes are largely shaped by existing gene pools rather than by the introduction of new resistance genes through fertilization [1,3,4]. The limited detection of isolate-derived genes in fertilizer metagenomes likely reflects methodological detection limits, emphasizing the complementary use of isolate genomics and metagenomics across abundance levels. For example, DNA extraction biases can skew representation toward more readily lysed taxa, while library preparation and sequencing steps, may influence detection of low-abundance genes, contributing to underrepresentation of certain taxa and functions [60–64].

At the bacterial community scale, fertilization altered resistome and mobilome without major shifts in dominant taxa or overall bacterial abundance, suggesting a decoupling between taxonomic composition and functional change. This pattern implies that responses of the resistome and mobilome are driven by shifts in ecological interactions rather than taxonomic turnover. Similar patterns have been observed in manure-amended soils [10], and animal-production settings, where increased ARG-MGE co-occurrence was observed without detectable shifts in community composition [12]. Such responses reflect the ability of soil microbiomes to maintain stability while remaining functionally flexible under environmental perturbations.

In this context, the mobilome was more responsive, showing coherent restructuring and strong associations with taxonomic (e.g. *Streptomyces*, and faecal-associated taxa) and functional (ARG abundance and diversity) gradients. Fertilizer origin further modulated these patterns: manure-based inputs were associated with MGEs linked to clinically relevant and opportunistic taxa, whereas compost-based inputs showed a wider range of associations with environmental and opportunistic groups. These differences suggest that BBF source materials define how the interactions of introduced genetic elements interact with soil communities. The observed coupling between ARGs and MGEs further supports the idea that these functional components respond together and MGEs act as a dynamic link that reshapes the resistome within the community. In contrast to the structured, largely immobile resistance observed in *Streptomyces* genomes, these patterns point to community-level processes shaped by the movement and rearrangement of existing genes rather than direct transfer of specific traits [5–7]. Despite these shifts, the overall changes were modest and mostly reversible, indicating a degree of resilience in the soil system [10].

Some limitations linked to our approach should be noted. Genomic analyses show potential functions but not actual activities of the bacterial species. Conclusively linking carbon metabolism, antibiotic production, and resistance would require integration of genomics with transcriptomic and metabolomic data. Metagenomics alone cannot fully capture horizontal gene transfer dynamics. For this, one would need to use approaches such as Hi-C or plasmid-resolved sequencing. Finally, longer-term experiments would be needed to assess whether repeated fertilization with the BBFs amplify or stabilize the patterns observed here.

Taken together, our results support our hypothesis that *Streptomyces* and BBFs impact soil resistomes through metabolically driven ecological processes shaped by the fertilizer composition and the soil community. Although the observed effects were modest and short-term, they are not negligible considering the evolutionary potential of the soil resistome, which may gradually alter resistance dynamics with long-term use of microbiologically active fertilizers.

## Supporting information

Supplementary Tablse S5

## Acknowledgements

We acknowledge the DNA Sequencing and Genomics Laboratory at the Institute of Biotechnology, University of Helsinki, supported by HiLIFE and Biocenter Finland, for sequencing services. T.-M.M. acknowledges travel support from the Luke doctoral programme. Computational resources were provided by CSC – IT Center for Science, Finland, through the Puhti computing environment.

## Supplementary material

Separate document

## Author contributions

**Taru-Marja Mäkinen**: Conceptualization, Data curation, Formal analysis, Investigation, Methodology, Project administration, Software, Visualization, Writing – original draft, and Writing – review and editing. **Melina Markkanen**: Methodology and Writing – review and editing. **Pietari Lahti-Nuuttila**: Investigation and Writing – review and editing. **Kirill Bodganov:** Investigation and Writing – review and editing. **Marko Virta**: Funding acquisition, Resources, and Writing – review and editing. **Jenni Hultman**: Conceptualization, Funding acquisition, Methodology, Project administration, Resources, Supervision, and Writing – review and editing. **Johanna Muurinen**: Conceptualization, Funding acquisition, Methodology, Project administration, Resources, Supervision, and Writing – review and editing.

## Conflicts of interest

The authors confirm no conflict of interest.

## Study funding

This work was supported by the Finnish Cultural Foundation to J.H. & J.M., the European Union’s Horizon 2020-funded project LEX4BIO to M.V. & J.M. (grant number 818309), which supported sample collection and metagenomic sequencing. The results reported in this paper reflect only the authors’ views, and the European Commission is not responsible for any use that may be made of the information it contains. This work was supported by the Research Council of Finland funding for the Multidisciplinary Center of Excellence in Antimicrobial Resistance Research to M.V. (364231 and 346125), which supported metagenomic sequencing and whole-genome sequencing. Additional support was provided by the Eemil Aaltonen Foundation to J.M., and T.-M.M. was supported by the RENEW Doctoral Programme at the University of Helsinki. Open access funded by Helsinki University Library.

## Data availability

All data supporting the findings of this study are included in the paper and as Supplementary Information. Adapter filtered metagenomic sequences are available in the European Nucleotide Archive (ENA) under accession number PRJEB114440. The isolate genome assemblies and filtered sequencing data are available in the ENA under accession number PRJEB67464. All scripts employed in this work are available from GitHub https://github.com/tarumarj/BBF_Streptomyces_project

## Use of AI Tools

Microsoft 365 Copilot (Microsoft, Redmond, WA, USA) was used to improve grammar, wording, and clarity of the manuscript text. Claude Sonnet 4.6 (Anthropic, San Francisco, CA, USA) was used to assist with code generation and refinement. All outputs were reviewed, edited, and validated by the authors. No AI tools were used to generate scientific conclusions or interpret results.

## Notes

https://github.com/tarumarj/BBF_Streptomyces_project

### Competing Interest Statement

The authors have declared no competing interest.

