## Supplementary Information for "Bio-based fertilizers shape soil microbiome, resistome and mobilome through metabolism of antibiotic-producing *Streptomyces*"

*Table S1, and S5 are available as separate Excell-documents

### Contents

#### Metagenomic data

**Supplementary tables…………………………………………………………………………… Pages 2–3**

**Table S1.** Per-sample metadata and community metrics (Sheet 1), genus-level microbial community composition as TSS-normalized relative abundances (Sheet 2), ARG abundances normalized by gene length and 16S rRNA gene counts (Sheet 3), MGE abundances normalized by gene length and 16S rRNA gene counts (Sheet 4), and BLASTn annotations of MGEs associated with the MGE NMDS ordination against the NCBI nt database (Sheet 5). Available as a separate Excel file (Table_S1.xlsx).

**Table S2** Pairwise PERMANOVA of bacterial communities beta diversity

**Table S3** Statistical comparisons of diversity (Shannon index) and normalized abundance across treatments.

**Table S4** Top 10 ARGs and MGEs contributors associated with ARG and MGE NMDS ordinations (envfit results)

#### Isolate data

**Supplementary tables……………………………………………………………… Separate document**

**Table S5** Supplementary genomic feature data for *Streptomyces* genomes. Sheet 1: sample metadata including genome assembly and annotation statistics, and sequence accessions. Sheet 2: ARG hits after BLASTn filtering (bitscore ≥100, query coverage ≥50%). Sheet 3: ARG presence/absence matrix. Sheet 4: BGC predictions after antiSMASH analysis and MIBiG annotation (similarity ≥15%). Sheet 5: CAZyme counts at the family level. Sheet 6: CAZyme counts at the class level. Sheet 7: MGE hits after BLASTn filtering (bitscore ≥50, query coverage ≥30%). Sheet 8: MGE presence/absence matrix. Sheet 9: MGE counts by element type. Available as a separate Excel file (Table_S5.xlsx).

**Supplementary Methods and Results…………………………………………………..…. Pages 3–6**

**Appendix S1** Detailed materials and methods for inhibition assays of *Streptomyces* isolates

**Appendix S2** Detailed functional gene analysis of *Streptomyces* genomes

**Supplementary figures……………………………………………………………………..…. Pages 6–9**

**Figure S1** Colony morphology of isolates grown on ISP7 or Gause's No. 1

**Figure S2** Spearman rank correlations between biosynthetic gene cluster (BGC) product types and CAZyme family abundances across 13 bacterial genomes.

**Figure S3** Cross-streak assay demonstrating antimicrobial activity of representative isolates against *Escherichia coli* DH5α.

**Supplementary information references……………………………………………………… Page 9**

**Table S2 Beta diversity analyses of bacterial communities**. Pairwise PERMANOVA using Bray–Curtis dissimilarities with 999 permutations were conducted for soil samples (n=4) only, against pre-fertilization soil ("Before), with Benjamini–Hochberg correction applied (p(BH)). DSP and PLP indicate fertilizer treatments; 6w denotes six weeks after fertilization.

| Category | Test | Comparison | R2 | F | df | p | p(BH) |
|---|---|---|---|---|---|---|---|
| Bacteria | PERMANOVA (pairwise) | Before vs DSP | 0.397 | 3.943 | | 0.022 | 0.022 |
| Bacteria | PERMANOVA (pairwise) | Before vs PLP | 0.309 | 2.678 | | 0.022 | 0.022 |
| Bacteria | PERMANOVA (pairwise) | Before vs 6w DSP | 0.491 | 5.797 | | 0.022 | 0.022 |
| Bacteria | PERMANOVA (pairwise) | Before vs 6w PLP | 0.478 | 5.505 | | 0.022 | 0.022 |

**Table S3 Statistical comparisons of diversity (Shannon index) and normalized abundance across treatments**. All comparisons are against pre-fertilization soil ("Before"). DSP and PLP indicate fertilizer treatments; 6w denotes six weeks after fertilization. n = 4 per group. P-values were adjusted using the Benjamini–Hochberg method (p(BH)).

| Category | Metric | Test | Comparison | Stat. | p | p(BH) |
|---|---|---|---|---|---|---|
| Bacteria | Shannon | Welch t | Before vs DSP | 5.075 | 0.009 | 0.036 |
| Bacteria | Shannon | Welch t | Before vs PLP | 1.474 | 0.233 | 0.233 |
| Bacteria | Shannon | Welch t | Before vs 6w DSP | -1.794 | 0.155 | 0.207 |
| Bacteria | Shannon | Welch t | Before vs 6w PLP | -2.754 | 0.058 | 0.115 |
| Bacteria | Abundance | Wilcoxon | Before vs DSP | 11 | 0.47 | 0.885 |
| Bacteria | Abundance | Wilcoxon | Before vs PLP | 10 | 0.665 | 0.885 |
| Bacteria | Abundance | Wilcoxon | Before vs 6w DSP | 7 | 0.885 | 0.885 |
| Bacteria | Abundance | Wilcoxon | Before vs 6w PLP | 11 | 0.47 | 0.885 |
| ARGs | Shannon | Welch t | Before vs DSP | 7.707 | 0.003 | 0.007 |
| ARGs | Shannon | Welch t | Before vs PLP | 5.127 | 0.003 | 0.007 |
| ARGs | Shannon | Welch t | Before vs 6w DSP | 6.492 | 0.006 | 0.008 |
| ARGs | Shannon | Welch t | Before vs 6w PLP | 1.309 | 0.246 | 0.246 |
| ARGs | Abundance | Wilcoxon | Before vs DSP | 16 | 0.03 | 0.041 |
| ARGs | Abundance | Wilcoxon | Before vs PLP | 16 | 0.03 | 0.041 |
| ARGs | Abundance | Wilcoxon | Before vs 6w DSP | 16 | 0.03 | 0.041 |
| ARGs | Abundance | Wilcoxon | Before vs 6w PLP | 12 | 0.312 | 0.312 |
| MGEs | Shannon | Welch t | Before vs DSP | 7.14 | 0.001 | 0.002 |
| MGEs | Shannon | Welch t | Before vs PLP | 6.151 | 0.001 | 0.002 |
| MGEs | Shannon | Welch t | Before vs 6w DSP | 4.086 | 0.007 | 0.009 |
| MGEs | Shannon | Welch t | Before vs 6w PLP | 3.735 | 0.012 | 0.012 |
| MGEs | Abundance | Wilcoxon | Before vs DSP | 16 | 0.03 | 0.061 |
| MGEs | Abundance | Wilcoxon | Before vs PLP | 16 | 0.03 | 0.061 |
| MGEs | Abundance | Wilcoxon | Before vs 6w DSP | 15 | 0.061 | 0.081 |
| MGEs | Abundance | Wilcoxon | Before vs 6w PLP | 12 | 0.312 | 0.312 |
| *Streptomyces* | Abundance | Wilcoxon | Before vs DSP | 2 | 0.11 | 0.22 |
| *Streptomyces* | Abundance | Wilcoxon | Before vs PLP | 0 | 0.03 | 0.12 |
| *Streptomyces* | Abundance | Wilcoxon | Before vs 6w DSP | 13 | 0.19 | 0.26 |
| *Streptomyces* | Abundance | Wilcoxon | Before vs 6w PLP | 12 | 0.31 | 0.31 |
| ARG-associated taxa | Abundance | Wilcoxon | Before vs DSP | 12 | 0.31 | 0.42 |
| ARG-associated taxa | Abundance | Wilcoxon | Before vs PLP | 16 | 0.03 | 0.12 |

| Category | Metric | Test | Comparison | Stat. | p | p(BH) |
|---|---|---|---|---|---|---|
| ARG-associated taxa | Abundance | Wilcoxon | Before vs 6w DSP | 4 | 0.32 | 0.42 |
| ARG-associated taxa | Abundance | Wilcoxon | Before vs 6w PLP | 9 | 0.89 | 0.89 |

**Table S4 Top 10 ARGs and MGEs contributors associated with ARG and MGE NMDS ordinations (envfit results)** P-values were adjusted using the Benjamini–Hochberg method (p(BH)). MGE BLASTn annotation results Supplementary table S1, sheet 5.

| Gene | NMDS1 | NMDS2 | r2 | p | p(BH) | Gene | NMDS1 | NMDS2 | r2 | p | p(BH) |
|---|---|---|---|---|---|---|---|---|---|---|---|
| aph(3'')-Ib_1 | -0.934 | 0.034 | 0.874 | 0.001 | 0.022 | 279_tnpA | -0.448 | 0.837 | 0.900 | 0.001 | 0.008 |
| tet(V)_1 | 0.589 | 0.639 | 0.755 | 0.001 | 0.022 | 710_tnpA | -0.481 | -0.818 | 0.900 | 0.001 | 0.008 |
| aph(6)-Id_1 | -0.709 | 0.428 | 0.685 | 0.001 | 0.022 | 1763_tnpA | -0.584 | 0.742 | 0.892 | 0.001 | 0.008 |
| CmlA9_1 | -0.803 | 0.103 | 0.655 | 0.001 | 0.022 | 501_tniA | -0.433 | -0.832 | 0.879 | 0.001 | 0.008 |
| dfrB3_2 | 0.288 | -0.750 | 0.646 | 0.001 | 0.022 | 1850_ISRj1 | 0.922 | -0.168 | 0.879 | 0.001 | 0.008 |
| tet(G)_3 | -0.775 | 0.120 | 0.614 | 0.001 | 0.022 | 391_tnpAcp2 | -0.355 | -0.853 | 0.853 | 0.001 | 0.008 |
| tet(G)_1 | -0.714 | -0.163 | 0.537 | 0.002 | 0.032 | 779_tnpA | -0.489 | 0.780 | 0.848 | 0.001 | 0.008 |
| ant(3'')-Ia_1 | -0.730 | 0.036 | 0.534 | 0.002 | 0.032 | 1440_istB | -0.430 | -0.814 | 0.848 | 0.001 | 0.008 |
| tet(X)_1 | -0.711 | -0.147 | 0.527 | 0.001 | 0.022 | 528_tnpA | -0.495 | 0.762 | 0.826 | 0.001 | 0.008 |
| erm(35)_1 | -0.707 | 0.035 | 0.502 | 0.001 | 0.022 | 2344_istA13 | -0.464 | -0.770 | 0.808 | 0.001 | 0.008 |

### Appendix S1. Detailed methodology for antimicrobial activity assays of *Streptomyces*

The isolation of bacterial strains, cultivation conditions, and sequencing-based taxonomic classification are described in the main Methods section. Briefly, eight isolates displaying *Streptomyces*-like morphology were obtained and classified based on taxonomic analysis. From these, four representative isolates (BBF42_03, BBF42_NS, BBF45_04, BBF45_11) were selected for antimicrobial assays to represent distinct species-level classifications within the isolate collection. *Kitasatospora aureofaciens* (prev. *Streptomyces*) strain HAMBI 51 (H-51) was obtained from the HAMBI culture collection (https://kotka.luomus.fi/culture/bac originating from Lederle Laboratories; ATCC12416a) [1] and used as a positive control in all growth and inhibition assays due to its known capability to produce antibiotics. *Escherichia coli* DH5α was used as the indicator strain.

International Streptomyces Project media ISP4, ISP5, and ISP7 were prepared following [2], with ISP7 prepared without glycerol. Isolates and the control strain were pre-cultivated on ISP7 plates at 28 °C for 4 days to obtain sufficient biomass for downstream assays. Antimicrobial activity was assessed using a modified cross-streak assay [3]. Each isolate and the control strain were streaked as a single central line onto ISP4 or ISP5 plates in duplicate and incubated at 28 °C for 7 days to allow extensive growth, sporulation, and production of secondary metabolites. The indicator strain *E. coli* DH5α was grown overnight at 37 °C on nutrient agar prior to use and streaked perpendicular to the central isolate streak using a sterile glass rod. Multiple streak passes were applied to generate variation in inoculum density across the plate. Plates were subsequently incubated at 37 °C for 2–3 days and monitored for bacterial growth.

Antimicrobial activity was evaluated visually based on inhibition of *E. coli* growth in relation to proximity to the isolate streak (Supplementary Fig. S4). On ISP4, metabolic activity of *E. coli* was inferred from a transition of the medium from opaque to translucent, whereas on ISP5 inhibition was assessed based on streak width, opacity, and the presence of discrete colonies. Reduced or absent growth near the isolate streak was interpreted as antimicrobial activity. Some isolates exhibited colony spreading beyond the original streak. In such cases, plates were excluded when spatial constraints prevented proper perpendicular streaking.

### Appendix S2 Detailed functional gene analysis of *Streptomyces* genomes

Functional gene profiles for antibiotic resistance genes (ARGs), antibiotic-production-associated biosynthetic gene clusters (BGCs), carbohydrate-active enzymes (CAZymes), and mobile genetic elements (MGEs) were compiled and curated as described in Materials and Methods (Supplementary Table S5, sheets 2–9). Across categories, retained hits were generally of moderate-to-high confidence, with ARGs and MGEs showing ≥70% sequence identity and varying query coverage, BGC similarities spanning a wide range reflecting both conserved and divergent secondary metabolite pathways, and CAZymes represented multiple families.

**Antibiotic resistance gene (ARG)** repertoires were strongly lineage-specific (Fig. 4; Supplementary Table S5, sheets 2–3). Only *ole(C)* was conserved across all genomes, occurring at moderate identity (74.3–89.9%) and broad query coverage (~69–98%), often in multiple genomic copies, suggesting a conserved but potentially multifunctional efflux-associated role. *Streptomyces speibonae* isolates (BBF42_03, BBF42_BW) displayed minimal and identical ARG profiles consisting of *aph(6)-Ia*, *blaFAR-1*, and *ole(C)*, all supported by moderate identity (~71–87%) and good coverage (~67–98%). In contrast, *S. diastaticus* isolate (BBF42_NS) encoded the most diverse ARG repertoire, including aminoglycoside resistance genes (*aac(6')*, *aph(3'')*), chloramphenicol resistance (*cmlV*), and macrolide-associated resistance (*tlr(C)*), with typical identities >70% and coverage ranging from ~51–93%. *S. cellulosae* (BBF45_07, BBF45_11, BBF45_15) isolates exhibited intermediate diversity, consistently encoding *tlr(C)*, *aph(6)-Ib*, *blaFAR-1*, and *ole(C)*, whereas the *Streptomyces* sp010548465 lineage (BBF45_04, BBF45_12) was restricted to *ole(C)* and aminoglycoside resistance genes. Reference genomes mirrored these trends, with *S. coelicolor* A3(2) encoding the most extensive repertoire, including high-confidence tetracycline (*tet*; 98.5% identity, 100% coverage) and erythromycin (*erm(O)*; 99.6% identity, 100% coverage) resistance genes.

**Antibiotic-production associated biosynthetic gene cluster (BGC)** profiles were similarly lineage-specific (Fig. 4; Supplementary Table S5, sheet 4). All predicted BGCs passing the ≥15% similarity threshold are reported in Supplementary Table S5, sheet 4; only BGCs with predicted antibiotic bioactivity based on MIBiG annotations were used in downstream analyses. The albaflavenone terpene cluster (BGC0000660) was universally conserved with 100% similarity. Beyond this shared component, BGC composition varied substantially in both class and similarity to reference clusters. *Streptomyces speibonae* isolates encoded a broad diversity of BGC classes, including nonribosomal peptide synthetase (NRPS) pathways, NRPS-independent (NI) siderophores, terpene biosynthetic clusters, and type III polyketide synthase (T3PKS) systems. Representative clusters included diazepinomicin (BGC0000679, 57% similarity), kinamycin (BGC0000236, 19% similarity), and aborycin (BGC0002285, 21% similarity), all associated with antibacterial activity. In addition, BBF42_03 uniquely encoded a partial daptomycin-associated NRPS cluster (BGC0000336, 25% similarity).

*Streptomyces diastaticus* (BBF42_NS) exhibited a distinct repertoire enriched in clinically relevant compounds, including NRPS-derived peptide antibiotics and type II polyketide synthase (T2PKS)-encoded aromatic polyketides. These included chloramphenicol (BGC0000893, 17% similarity), echinomycin (BGC0000339, 88% similarity), and enterocin (BGC0000220, 95% similarity), reflecting a combination of conserved and divergent biosynthetic pathways with antibacterial, antifungal, and cytotoxic activities. *S. cellulosae* isolates displayed reduced BGC diversity, dominated by ribosomally synthesised and post-translationally modified peptide (RiPP)-associated pathways and T2PKS-derived clusters. These included aborycin (BGC0002285, ~28% similarity) and alnumycin-related clusters (BGC0000195, 71% similarity), often associated with azole-containing RiPP features. Large NRPS and type I polyketide synthase (T1PKS)-based antibiotic pathways were consistently absent from this lineage.

In contrast, the *Streptomyces* sp010548465 lineage encoded broader and more distinctive BGC repertoires, including hybrid NRPS–polyketide pathways such as cinnapeptin (NRPS/highly reducing T2PKS; BGC0002108, ~28% similarity), furan-associated methylenomycin A (BGC0000914, ~23% similarity), and the terpene-derived cyclooctatin pathway (BGC0000677, 100% similarity). Across all lineages, similarity values ranged from near-identical matches (≥90–100%) to low similarity (~15–30%), indicating a mixture of conserved biosynthetic pathways and potentially novel or divergent secondary metabolite variants with ecological and competitive relevance. Overall, BGC profiles were consistent with broad biosynthetic potential across lineages, with both shared and lineage-specific clusters suggestive of niche-related differences in secondary metabolism.

**Carbohydrate-active enzymes (CAZymes)** exhibited the most pronounced functional differentiation among lineages and were strongly indicative of variation in the capacity to degrade complex carbon substrates (Fig. 4; Supplementary Table S5, sheets 5–6). Differences in total CAZyme content were primarily driven by variation in glycoside hydrolase (GH) and carbohydrate-binding module (CBM) families, with additional contributions from auxiliary activity (AA) enzymes, carbohydrate esterases (CEs), glycosyltransferases (GTs), and polysaccharide lyases (PLs).

GH families directly reflect the ability to hydrolyze recalcitrant polysaccharides. Families associated with cellulose degradation (GH5, GH6, GH9, GH48) and hemicellulose breakdown (GH10, GH11, GH26) were most abundant in *S. speibonae* and *S. cellulosae*, supporting efficient utilization of lignocellulosic plant material. These lineages exhibited high overall GH richness (128 in *S. speibonae* and 133–134 in *S. cellulosae*), consistent with broad substrate utilization. In contrast, *S. diastaticus* showed a reduced GH repertoire (GH = 83), lacking several hemicellulose-active families, indicating more limited capacity to degrade complex plant-derived substrates. The *Streptomyces* sp010548465 lineage showed intermediate GH diversity (101–103), while *S. coelicolor* A3(2) exhibited the highest overall GH richness (179), including additional specialized families. CBMs, which facilitate enzyme binding to insoluble substrates, were enriched in lineages with high degradative capacity, particularly CBM2, CBM13, and CBM48 associated with cellulose and hemicellulose targeting. Their expansion in *S. speibonae* and *S. cellulosae* supports efficient enzymatic access to complex plant polymers, whereas reduced CBM representation in *S. diastaticus* paralleled its reduced GH complement.

The remaining CAZyme classes showed consistent patterns across lineages. AA enzymes (e.g. AA1, AA2, AA3, AA10), which mediate oxidative breakdown of lignin, were most abundant in *S. speibonae* and *S. cellulosae*, supporting lignocellulose degradation through combined oxidative and hydrolytic processes. CE families (e.g. CE4, CE14) involved in deacetylation of hemicellulose and pectin were similarly enriched in these lineages, facilitating access to otherwise protected polysaccharide backbones. PLs, responsible for cleavage of uronic acid-containing polysaccharides such as pectin, were absent in *S. diastaticus* but present at low abundance in other isolates, indicating lineage-specific capacity for pectin degradation. Overall, CAZyme profiles reveal clear ecological differentiation, with *S. speibonae* and *S. cellulosae* specialized for degradation of complex lignocellulosic substrates, *S. diastaticus* showing reduced and potentially more specialized carbon utilization, and *S. coelicolor* exhibiting broad metabolic versatility.

**Mobile genetic elements** (MGEs) showed limited diversity but substantial copy number variation (Fig. 4; Supplementary Table S5, sheets 7–9). Only ISCau1 and transposase-associated *tnpA* variants were consistently detected. ISCau1 (accession NC012590) occurred in all genomes as a partial sequence with moderate identity (74.2–76.1%) and low but consistent coverage (~43–45%), indicative of fragmented or inactive elements. A total of 14 *tnpA* variants were identified, typically showing moderate identity (~70–90%)

and variable coverage (~30–100%). Two variants dominated: 1640_*tnpA* (detected in 10/13 isolates, 1–9 copies per genome, accession AF099014.1, *Streptomyces coelicolor* A3(2) transposase), and 1764_*tnpA* (7/13 isolates, 1–6 copies, accession X97015.1, *Paenarthrobacter nicotinovorans tnpA* gene), together comprising the majority of MGE content. Total MGE abundance ranged from 1–7 elements per genome, with highest variability observed in *S. cellulosae* (3–7 elements) and lowest counts in the *Streptomyces* sp010548465 lineage (1–4 elements). The reference genome *S. coelicolor* A3(2) carried the highest MGE burden, including multiple near-identical *tnpA* copies (99–100% identity). DSP and PLP isolates did not differ markedly in overall MGE load, suggesting that transposable element content is more strongly associated with lineage identity than with sampling site. The low MGE diversity across all isolates, dominated by *tnpA*-associated elements, is consistent with the general pattern observed in *Streptomyces* genomes, where IS elements are present but horizontal gene transfer via MGEs appears less frequent than in many other bacteria.

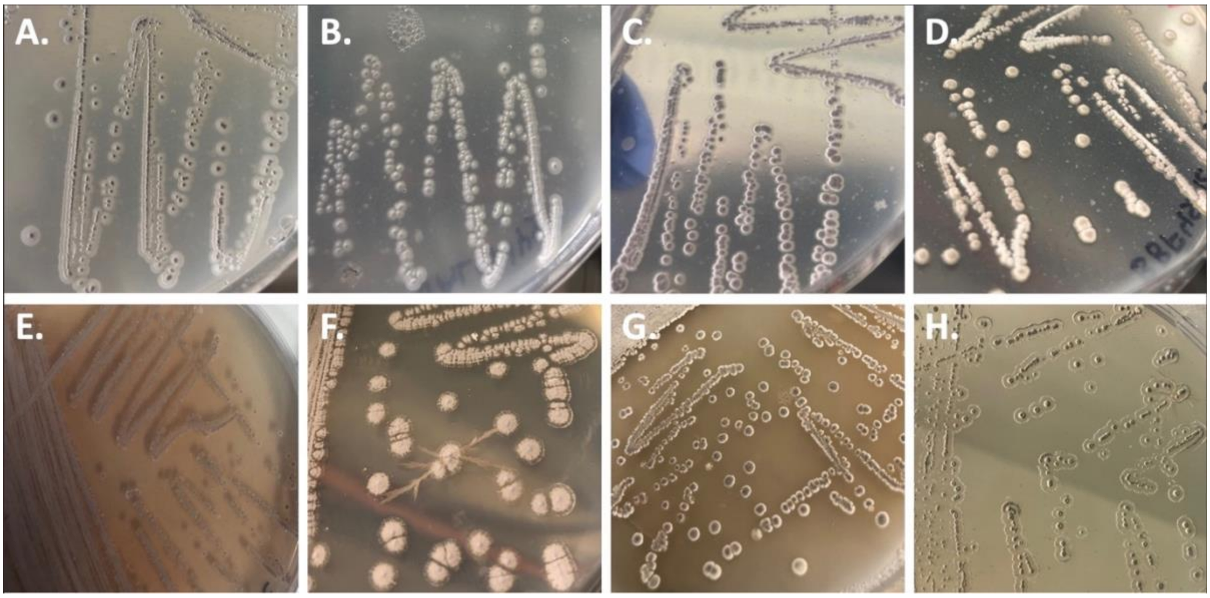


**Figure S1 Colony morphology of the isolates.** Colonies were generally dry, circular to punctiform, and varied in margin and elevation depending on the medium. On Gause's No. 1, colonies exhibited smooth edges and raised to umbonate elevations, with well-developed aerial and substrate mycelium ranging from white to grey and pale yellow to brownish in substrate coloration (A–D, G). On ISP-7-gly, colonies showed rough to filamentous margins, flatter to umbonate elevations, and in some cases powdery aerial mycelium and diffusible brown pigment (E, F, H). Representative isolates: BBF42_03 (A), BBF42_BW (B), BBF45_04 (C), BBF45_12 (D), BBF45_07 (E), BBF45_11 (F), BBF45_15 (G), and BBF42_NS (H).

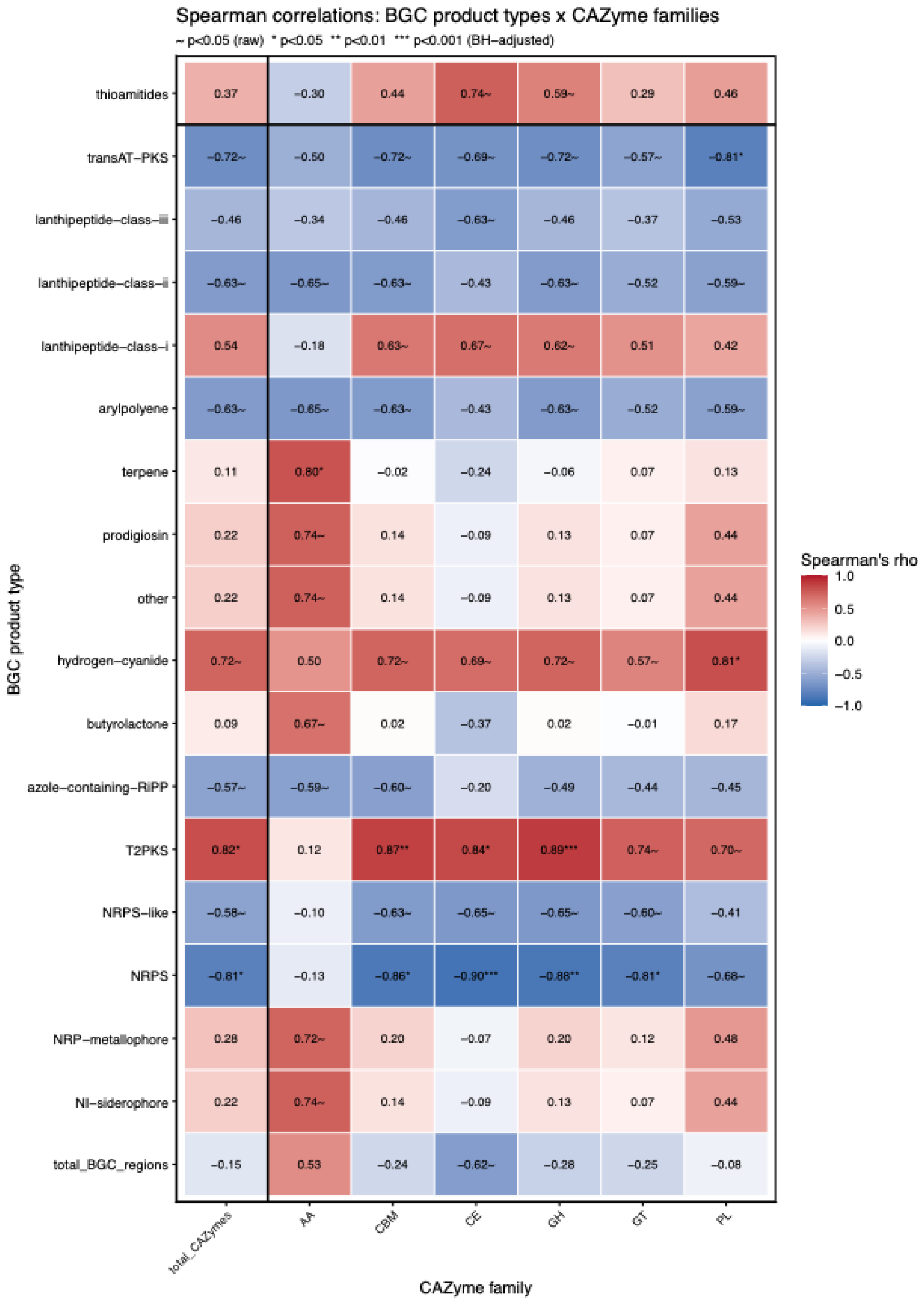


**Figure S2 pearman rank correlations between biosynthetic gene cluster (BGC) product types and carbohydrate-active enzyme (CAZyme) family abundances across 13 Streptomyces isolates.** Heatmap cells show Spearman's ρ for each BGC product type (rows) × CAZyme family (columns) pair. Colour intensity reflects the strength and direction of the correlation, with red indicating positive and blue indicating negative correlations. Only BGC product types and CAZyme families with at least one nominally significant correlation (p < 0.05, uncorrected) are shown; shown BGC product types are: nonribosomal peptide synthetase (NRPS), NRPS-like, type II polyketide synthase (T2PKS), trans-acyltransferase polyketide synthase (transAT-PKS), hydrogen cyanide (hydrogen-cyanide), arylpolyene, lanthipeptide class I–III (lanthipeptide-class-i/ii/iii), azole-containing ribosomally synthesised and post-translationally modified peptide (azole-containing-RiPP), NI-siderophore, nonribosomal peptide (NRP)-metallophore, butyrolactone, prodigiosin, thioamitides, and total BGC regions. CAZyme families shown are: auxiliary activities (AA), carbohydrate-binding modules (CBM), carbohydrate esterases (CE), glycoside hydrolases (GH), glycosyltransferases (GT), polysaccharide lyases (PL), and total CAZymes. Cells marked ~ are nominally significant (p < 0.05, uncorrected) but did not survive Benjamini–Hochberg (BH) false discovery rate correction. The bold vertical and horizontal lines separate total counts (total CAZymes, total BGC regions) from individual families and product types, respectively. Significance thresholds after BH correction: * p < 0.05, ** p < 0.01, *** p < 0.001.

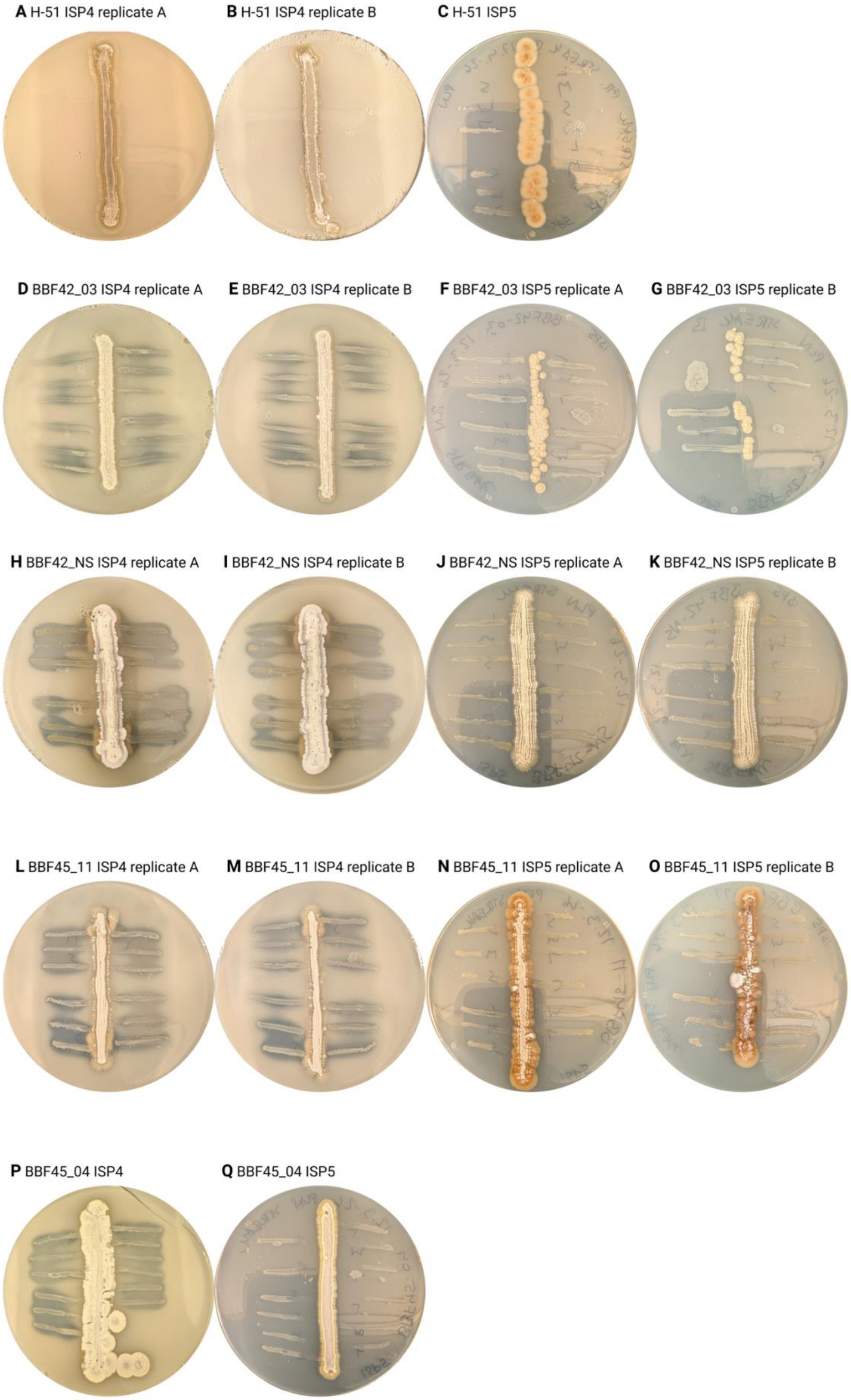
A H-51 ISP4 replicate A
B H-51 ISP4 replicate B
C H-51 ISP5
D BBF42_03 ISP4 replicate A
E BBF42_03 ISP4 replicate B
F BBF42_03 ISP5 replicate A
G BBF42_03 ISP5 replicate B
H BBF42_NS ISP4 replicate A
I BBF42_NS ISP4 replicate B
J BBF42_NS ISP5 replicate A
K BBF42_NS ISP5 replicate B
L BBF45_11 ISP4 replicate A
M BBF45_11 ISP4 replicate B
N BBF45_11 ISP5 replicate A
O BBF45_11 ISP5 replicate B
P BBF45_04 ISP4
Q BBF45_04 ISP5

**Figure S3 Cross-streak assay demonstrating antimicrobial activity of representative isolates against *Escherichia coli* DH5α**. Antimicrobial activity was assessed using a modified cross-streak method on ISP4 and ISP5 media. The central vertical streak corresponds to the *Streptomyces* isolates (D–Q) or the positive control strain *Kitasatospora aureofaciens* (formerly *Streptomyces*) HAMBI 51 (H-51) (A–C), while perpendicular streaks represent *E. coli* DH5α applied at varying inoculum densities. The four isolates shown (BBF42_03, BBF42_NS, BBF45_04, BBF45_11) represent distinct species-level taxonomic groups. Biological replicates (A, B) are included where available. In replicate assays, BBF45_04 formed colonies outside the original streak, and H-51 showed growth only on one ISP5 plate.

All isolates inhibited *E. coli* growth in proximity to the central streak to varying degrees, with inhibition strongest closest to the producer strain. The density of *E. coli* inoculum did not appear to substantially affect the observed inhibition. Overall, inhibition was more pronounced on ISP4 medium. The positive control (H-51) fully inhibited *E. coli* on ISP4 (A, B) and partially on ISP5 (C). Isolates BBF42_03 (D–G) and BBF42_NS (H–K) exhibited slightly stronger inhibitory activity than BBF45_04 (P, Q) and BBF45_11 (L–O). Variability between replicates was generally minor but occasionally observed, likely reflecting differences in colony growth or medium consistency. Due to variation in medium appearance, precise quantification of inhibition was not feasible.
